# Viral lineage and mode of exposure modulate within host spatial dynamics of influenza A viruses

**DOI:** 10.1101/2025.11.03.686270

**Authors:** Christina M. Leyson, Nahara Vargas-Maldonado, Matthew Gaddy, Vedhika Raghunathan, Lucas M. Ferreri, Meher Sethi, Kayle Patatanian, Silvia Carnaccini, Ketaki Ganti, David VanInsberghe, Anice C. Lowen

## Abstract

The upper and lower respiratory tracts (URT and LRT) present distinct environments for influenza A virus (IAV) replication. Their differential features have major implications for viral evolutionary dynamics, transmission potential, and pathogenesis. To investigate the implications of differential viral replication in the URT and LRT, we assessed dispersal of IAVs throughout the guinea pig respiratory system. Guinea pigs were inoculated intranasally with a 300 μL volume to deliver inoculum to both the URT and LRT. Two strains were used to represent the circulating seasonal IAV lineages: influenza A/TX/50/2012 (H3N2) and influenza A/CA/07/2009 (H1N1) virus. The inclusion of a diverse genetic barcode enabled high-resolution tracing of viral dispersal for the H1N1 virus. While infectious virus was consistently detected in the URT, the H1N1 virus could be detected in LRT while the H3N2 virus could not. To determine whether replication of the H1N1 virus in the LRT extends to other modes of infection, virus distribution was evaluated following infection via aerosol exposure or transmission. Infectious virus in lung homogenates was observed in both cases, confirming the LRT tropism of the H1N1 virus. Sequencing genetic barcodes revealed that diversity was largely maintained in nasal samples and trachea but contracted upon dispersal to the lungs. This loss of diversity was associated with increased distance to and branching from the major airways, implicating long distance dispersal through the airways in imposing within-host population bottlenecks. These data underline the implications for within-host viral dynamics of the distinct environments of the upper and lower respiratory tracts.

**Importance:** The upper (URT) and lower (LRT) respiratory tracts create different conditions for influenza A virus (IAV) spread and evolution. We studied how the virus moves through guinea pigs’ airways after infection with H3N2 or H1N1 strains of IAV. Whether delivered intranasally, by aerosol or by transmission, the H1N1 virus replicated in the nasal cavity, trachea, and lungs. By contrast, the H3N2 virus stayed mostly in the nasal cavity. Genetic barcodes were used to track how the H1N1 virus moved and changed. The populations replicating in the nasal cavity and trachea maintained high diversity but those sampled from the lungs showed low diversity. This bottlenecking effect was stronger for viral populations present deeper in the lungs. These findings show that the different environments of the URT and LRT strongly shape how influenza spreads and evolves inside a host.

## Introduction

Influenza A viruses (IAVs) are known to infect many cell types within their hosts and can propagate within the upper and lower respiratory tracts of mammals (1–3). The upper respiratory tract (URT) comprises the tissues of the nose, nasal cavity, mouth, pharynx, and larynx, and the lower respiratory tract (LRT) is composed of the trachea and lungs. These regions of the respiratory tract differ markedly in the cell types present and the frequency and spatial arrangements of those cell types (2, 4–6). In turn, the URT and LRT are characterized by differing levels of mucus, ciliary action and immune reactivity (2, 4). In addition, these regions differ in their temperature and the types and availability of sialic acid receptors used by IAVs (7, 8). These features are all expected to shape the tropism of IAVs for URT and LRT tissues and the characteristics of viral dispersal within each region.

The dynamics of IAV infection in the upper and lower respiratory tracts are important to understand. In the clinic, infections of the LRT are especially consequential and are a leading cause of morbidity and mortality (9–11). Infections of the URT, on the other hand, are important for transmission and therefore viral evolution at the host population level (12–15). In a ferret model, we and others have shown with H1N1 IAV from the 2009 pandemic that viral populations in the URT are well mixed and subject to relatively little stochastic loss of diversity over time (16–18). In contrast, populations in the LRT are comprised of multiple spatially distinct, low diversity subpopulations that are dissimilar in their composition (16–18).

In this study, we sought to extend investigation of IAV dynamics within the respiratory tract to a guinea pig model, given the utility of this model host for studying IAV transmission and evolutionary dynamics (19–29). We therefore evaluated the replicative potential of both pandemic H1N1 (pH1N1) and seasonal H3N2 viruses in the guinea pig URT and LRT. Since the pH1N1 strain showed broader tropism, we used this system to compare the spatial distribution of viral populations seeded through three different modes of infection and to evaluate population genetic structure using a virus carrying a diverse genetic barcode.

## Results

### Destination of intranasal inoculum in the respiratory tract

Intranasal delivery is commonly used to inoculate animals with respiratory viruses. To evaluate how the destination of a liquid inoculum varies with volume, we administered 30, 100, or 300 *µ*L of blue tissue marking dye intranasally to guinea pigs and then harvested respiratory tissues for gross examination. We observed that the nasal passages were stained in all volumes tested (Figure 1). However, staining in the oral cavity and LRT, including trachea and lungs, were only observed with the highest volume (300 *µ*L). We also noted that the esophagi of all three guinea pigs receiving 300 *µ*L were stained, indicating that some of the 300 *µ*L inoculum is swallowed.

**Figure 1.**
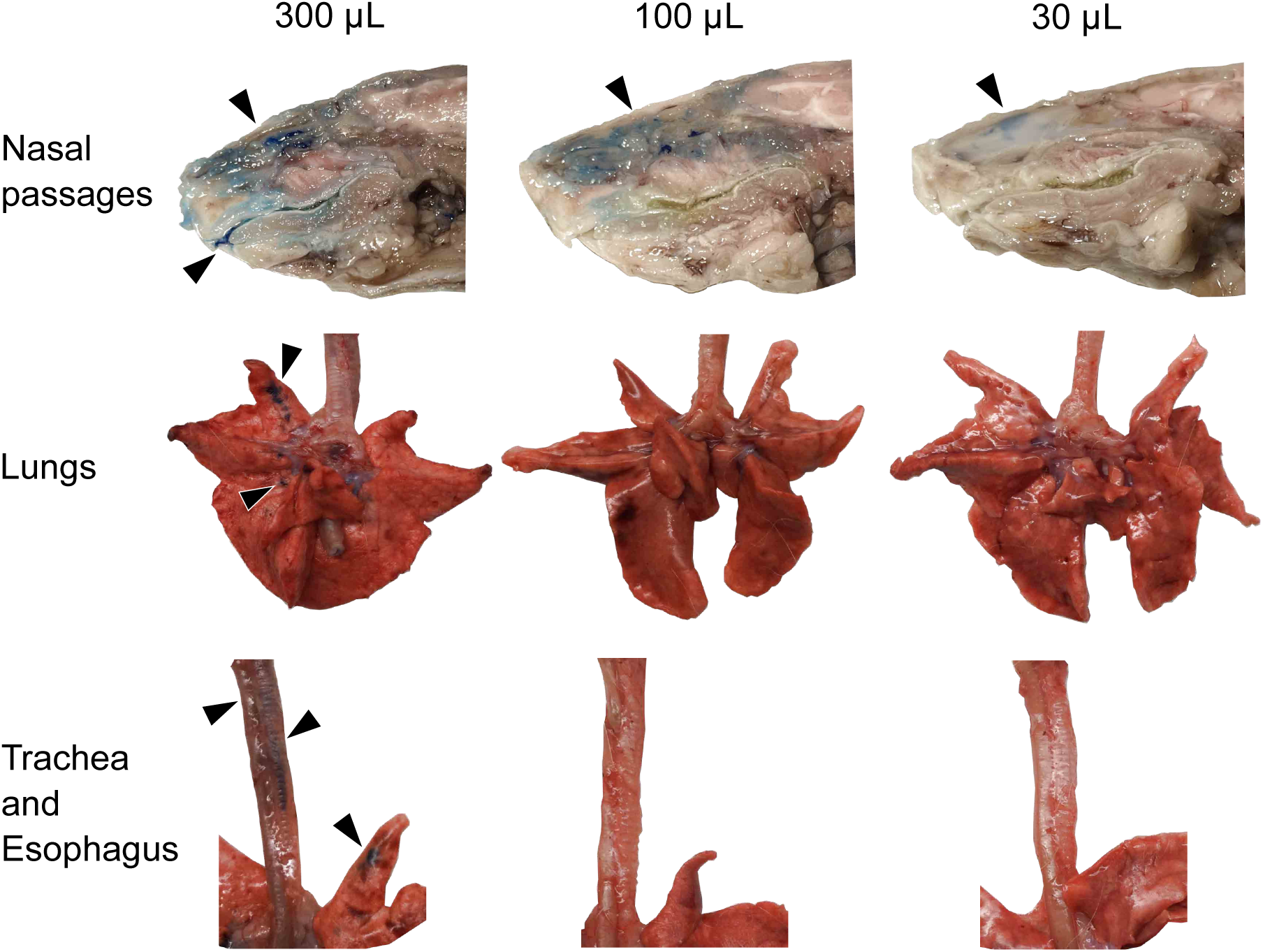
A 300 μL inoculum reaches the lungs of guinea pigs whereas 30 μL and 100 μL inocula remain in the nasal cavity. The volume of the tissue marking dye administered is indicated across the top of the figure. Anatomical structures are labeled on the left. Black arrows indicate tissue dye markings.

### pH1N1 replicates in the LRT whereas seasonal H3N2 does not

The extent of LRT virus replication during influenza virus infections is unclear, especially in the guinea pig model. To address this knowledge gap, we inoculated guinea pigs intranasally with either influenza A/Texas/50/2012 (H3N2) virus, a seasonal strain, or influenza A/California/07/2009 (H1N1) virus, a pandemic strain, using a 300 *µ*L inoculum. This high volume was used to ensure that the inoculum would reach the lungs (Figure 1). We collected nasal lavage from one group of guinea pigs and respiratory tissues from a distinct group to avoid artificial virus dispersal during nasal lavage. Both H3N2 and pH1N1 viruses were found at high titers in nasal lavage (Figure 2A). The peak of viral replication at this site for both viruses was at 2 days post-inoculation (dpi). In contrast to nasal lavages, infectious virus in the lungs was detected only in animals inoculated with pH1N1 (Figure 2C). Furthermore, the peak of viral positivity in LRT was at 1 dpi, suggesting that virus replication kinetics in the LRT are faster compared to the URT. Immunohistochemistry for influenza A nucleoprotein corroborated the difference in LRT replication between seasonal H3N2 and pH1N1: while viral antigen was abundant in the nasal epithelia of guinea pigs infected with either virus, viral antigen in the lung epithelium was observed in the major airways only of animals inoculated with pH1N1 (Figure 2D, Table 1). The profile of antigen positive cells in the nasal passages was similar between animals inoculated with pH1N1 or H3N2, comprising respiratory and olfactory epithelial cells, olfactory neurons, goblet cells, and leukocytes. Catarrhal exudates in the nasal passages were also antigen positive. In the lung, antigen positivity was observed in the bronchiolar mucosal epithelium, pneumocytes, and leukocytes.

**Figure 2.**
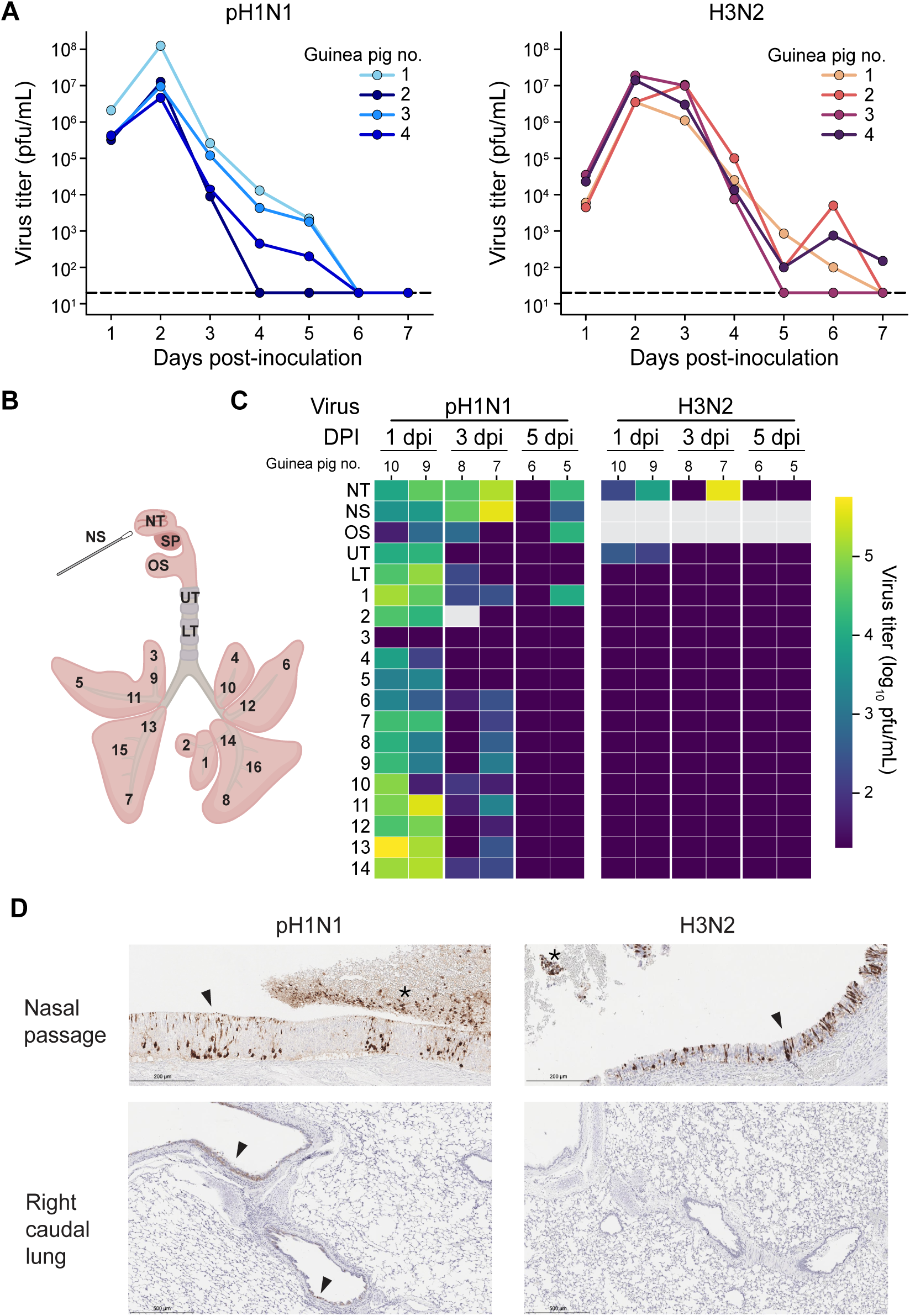
Both pH1N1 and H3N2 viruses replicate well in the URT but only pH1N1 replicates in the lungs. (A) Viral titers in nasal washes. Dashed lines indicate the limit of detection (50 pfu). (B) Schematic of the respiratory tissues sampled. NS: nasal swab, NT: nasal turbinates, SP: soft palate, UT: upper trachea, LT: lower trachea, 1-16: lung lobes. (C) Heat map showing viral titers according to the tissue locations labeled in (B). (D) Immunohistochemical staining for influenza A virus nucleoprotein is visible as dark brown coloring. Arrows indicate antigen positive cells within intact epithelia. Asterisks mark positive catarrhal exudate. DPI = day post-inoculation.

**Table 1.** Immunohistochemistry scores for influenza A nucleoprotein in guinea pigs infected with pH1N1 virus via different routes^1^.

| Route of exposure<br>Timepoint<br>Guinea pig ID# | pH1N1 |  |  |  |  |  | H3N2 |  |
| --- | --- | --- | --- | --- | --- | --- | --- | --- |
|  | intranasal |  | aerosol |  | recipient |  | intranasal |  |
|  | 3 dpi<sup>2</sup> |  | 3 dpi |  | 5 dpi |  | 3 dpi |  |
|  | 1-12 | 1-5 | 6-1 | 6-2 | 2-16 | 5-22 | 1-13 | 1-14 |
| Nasal passage | 4 | 3 / 5 / 5 | 0 / 2 / 3 | 0 / 0 / 0 | 0 / 2 / 3 | 0 / 1 / 2 | 0 / 3 / 3 | 0 / 2 / 2 |
| Trachea | 0 / 0<sup>1</sup> | 0 / 0 | 0 / 0 | 0 / 0 | 0 / 0 | 0 / 0 | 0 / 0 | 0 / 0 |
| Left caudal lung | 1 | 1 | 0 | 0 | 0 | 0 | 0 | 0 |
| Left middle lung | 1 | 1 | 0 | 1 | 0 | 0 | 0 | 0 |
| Right caudal lung | 1 | 2 | 0 | 0 | 0 | 0 | 0 | 1 |
| Right middle lung | 1 | 2 | 0 | 0 | 0 | 0 | 0 | 0 |

<sup>1</sup> Scores were given 0 for no observed staining, up to 5 with >60% positivity.
<sup>2</sup> DPI = days post-inoculation. For recipients, the dpi refers to the timing of infection of the donor that the animal was paired with.
<sup>3</sup> If multiple sections of a tissue were taken, each section was scored separately and results are separated by /.

### LRT replication of pH1N1 is also observed after aerosol exposure and transmission

To extend our characterization of IAV tropism in the guinea pig to additional exposure contexts, we examined the spatial distribution of pH1N1 replication following infection via an aerosol route or through co-housing with an infected animal (transmission). For aerosol exposures, we aerosolized either 5×10^4^ pfu (high dose) or 5×10^2^ pfu (low dose) per guinea pig in particles of approximately 1-5 *µ*m diameter. We measured the uptake of virus in tissues 1 h after inoculation by quantitative RT-PCR and estimate that 1.53 x 10^2^ pfu (high dose) or 1.53 pfu (low dose) was successfully delivered to the respiratory tissues. Inoculation with the higher dose led to productive infection in guinea pigs that was detectable in the nasal washes (Figure 3A) and respiratory tissues (Figure 3B) throughout the study period. Inoculation with the lower dose, by contrast, did not yield detectable virus in nasal washes (Figure 3A) but did result in low-level viral detection at 2 dpi in 2-3 lung tissue sections per guinea pig (Figure 3B). Viral antigen was abundantly found in the nasal passages whereas few viral antigen positive cells were found in the lungs (Figure 3C, Table 1), consistent with virus titers observed. The virus positivity observed in the lower dose group suggests limited viral replication initiated by virus-laden aerosols inhaled directly to lung tissue.

**Figure 3.**
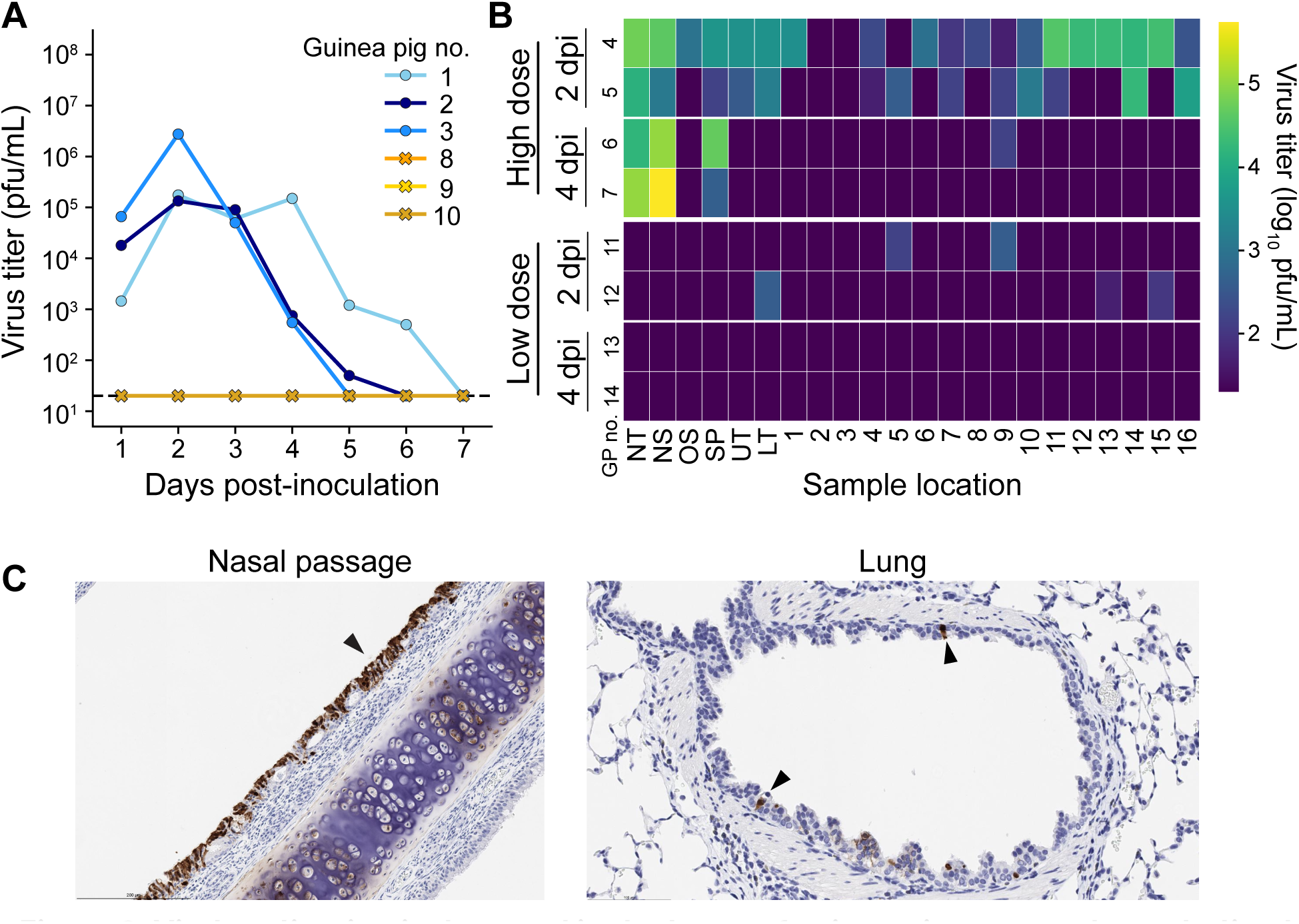
Viral replication is detected in the lungs of guinea pigs exposed to nebulized pH1N1 virus. Guinea pigs 1-7 received 1.53×10^2^pfu and guinea pigs 8-14 received 1.53×10^0^pfu. (A) Viral titers in nasal washes. Dashed lines indicate the limit of detection (50 pfu). (B) Heat map showing viral titers according to the tissue locations labeled in Figure 1(B). DPI = day post-inoculation. NS: nasal swab, NT: nasal turbinates, SP: soft palate, UT: upper trachea, LT: lower trachea, 1-16: lung lobes. (C) Immunohistochemical staining for influenza A virus nucleoprotein at day 2 post-inoculation. Black arrows indicate antigen positive cells with dark brown color.

Transmission experiments were carried out by co-housing a recipient guinea pig with a donor that was intranasally inoculated with either 5×10^4^ (first experiment) or 1×10^4^ pfu (second experiment) of pH1N1 virus 24 h previously. These two experiments yielded 1/3 and 2/3 transmission events, respectively, giving a total of three recipient animals for analysis (Figure 4 A). In nasal washes, recipient guinea pigs had peak viral titers at day 5-6 post-donor inoculation. Viral load in URT tissue sections, and particularly in nasal turbinates, was high with titers up to 10^5^ pfu/mL at day 4 or 5. In contrast, viral positivity in lung sections of transmission recipients was sparse and low in magnitude relative to that seen following intranasal inoculation or aerosol exposure (Figure 4B).

**Figure 4.**
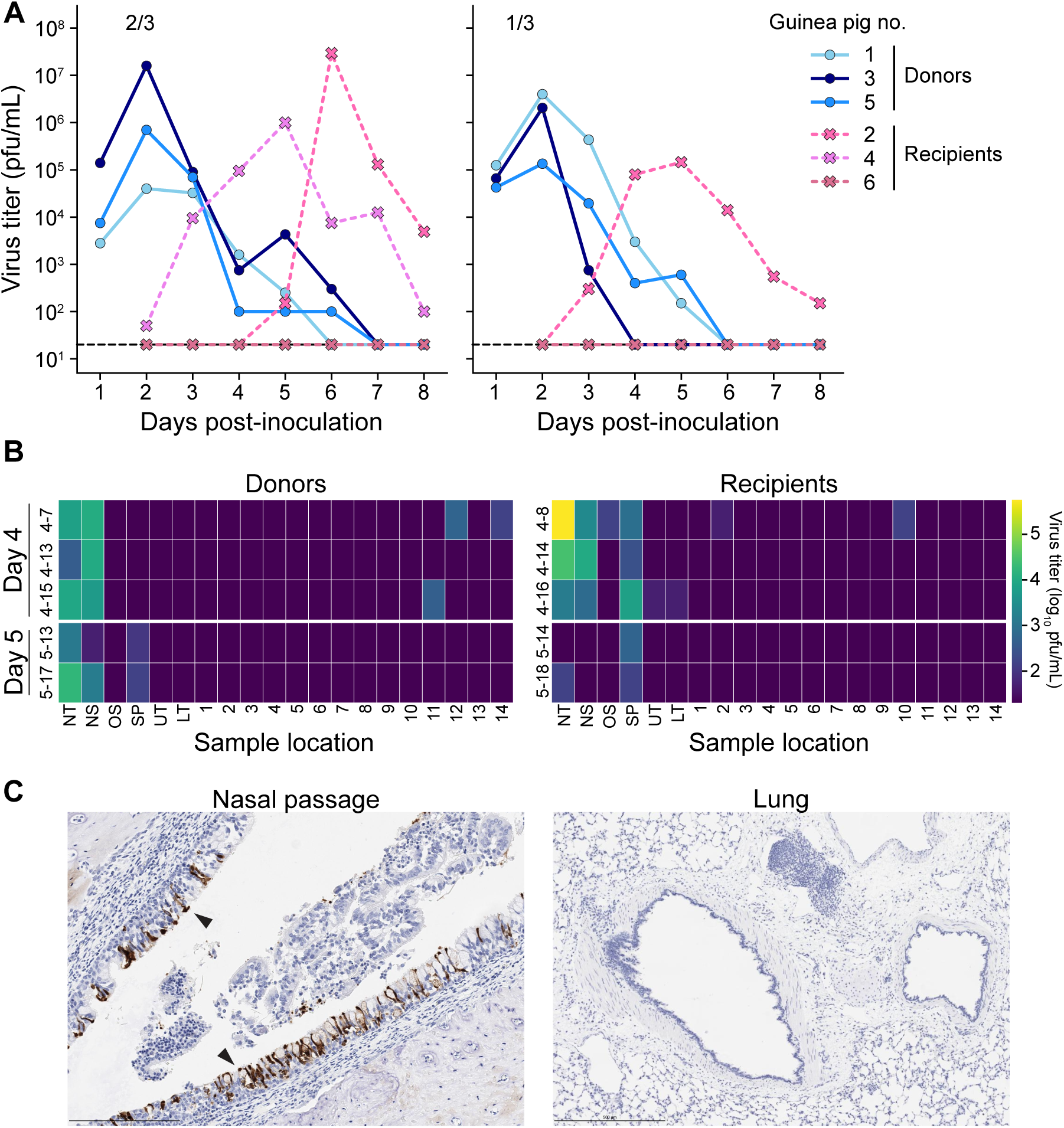
Transmission recipient guinea pigs show robust replication of pH1N1 virus in the URT but limited replication in the LRT. (A) Viral titers in nasal washes. Dashed lines indicate the limit of detection (50 pfu). (B) Heat map showing viral titers according to the tissue locations labeled in Figure 1(B). DPI = day post-inoculation. NS: nasal swab, NT: nasal turbinates, SP: soft palate, UT: upper trachea, LT: lower trachea, 1-16: lung lobes. The donor-recipient pairs share the same row. (C) Immunohistochemical staining for influenza A virus nucleoprotein at day 4 post-exposure. Black arrows indicate antigen positive cells with dark brown color.

Immunohistochemistry for influenza A virus antigen showed similar trends, with much higher antigen positivity in the nasal passages compared to the lungs and trachea (Figure 4C, Table 1). The cell types positive for influenza A virus antigen were similar to those observed for intranasal inoculation.

### Viral diversity is maintained in the URT and reduced upon dispersal to the LRT

Determination of viral titers and antigen positivity at multiple tissue sites gives valuable insight into viral tropism, but only a high-level view of how the within-host viral population changes over space and time. To obtain greater resolution on viral population dynamics, we used a previously established viral barcoding system (16, 30). In all of the experiments described, the inoculum used to infect animals with pH1N1 virus was a barcoded library of viruses (16). This library comprises approximately 4,000 genetic lineages that differ from each other by 1–12 synonymous mutations that are clustered within a 100-nucleotide region of the NA gene segment (the barcode region). Deep sequencing of this region in virological samples collected from various regions of the respiratory tract allows for high-resolution mapping of viral lineages as they disperse within a host.

Sequencing of viral barcodes in nasal washes revealed that high diversity was maintained throughout the course of infection for all four guinea pigs inoculated intranasally with pH1N1 virus (Figure 5A). Examination of viral barcodes in swabs and tissue sections showed that viral diversity was likewise maintained in nasal swabs, nasal turbinates, oral swabs, and tracheal tissues (Figure 5B-C). In contrast, viral populations sampled from the lung tissues showed a broad range of barcode diversities, with diversity similar to that seen in the URT in some locations and a single barcode comprising >90% of the population in several locations (Figure 5B). Some anatomical patterns were apparent in the diversity profiles seen in the LRT (Figure 5C): The lung lobes in the caudal (lower) region had significantly higher diversity compared to those in the apical (upper) region. Within each lung lobe, the sections proximal to the major airways also had significantly higher diversity compared to those distal to the main airways. No differences in viral diversity were observed between the left and right lung lobes.

**Figure 5.**
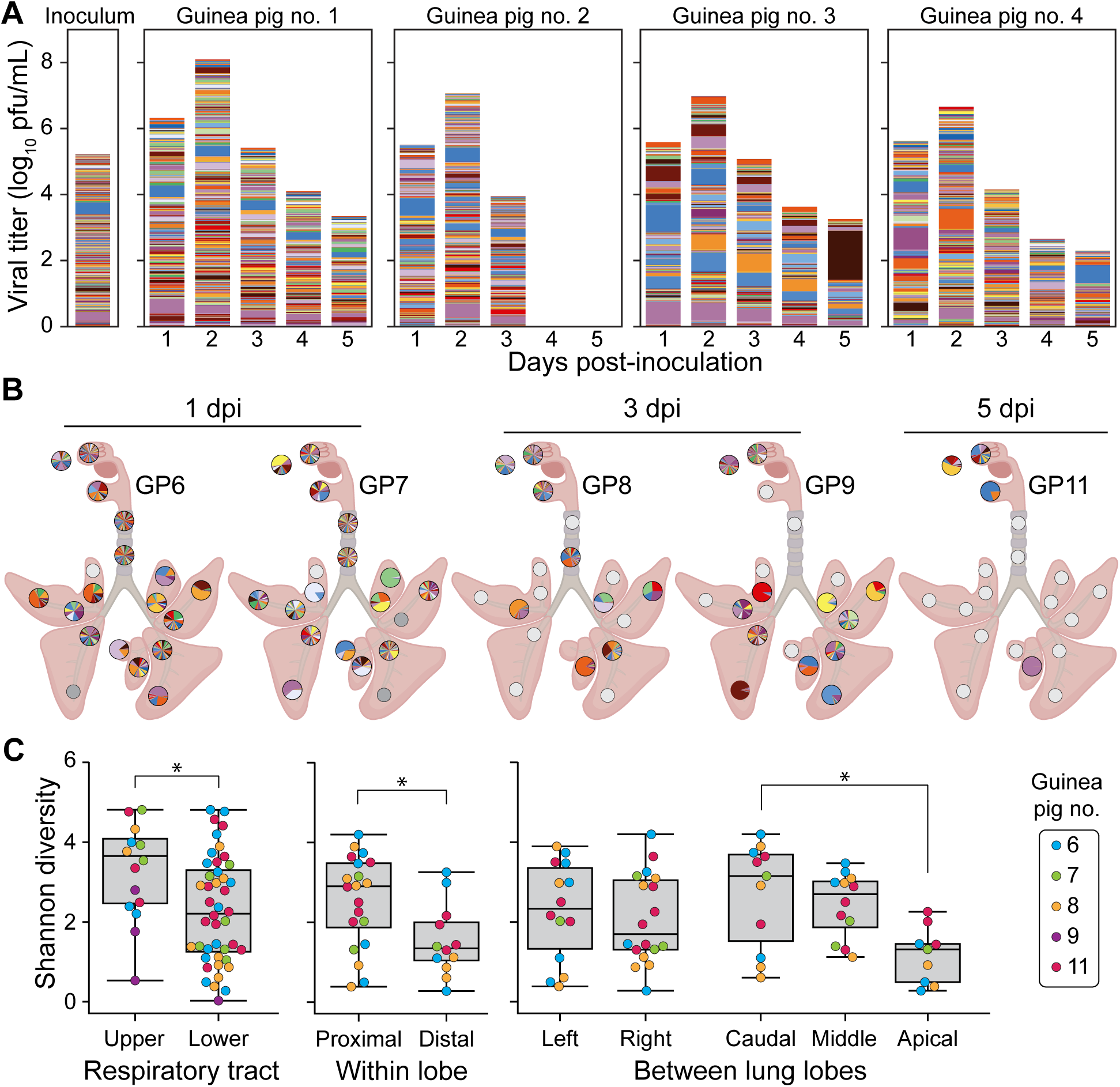
Barcode diversity is maintained in the URT but declines with increased distance from and branching of the main airways. Viral barcodes detected in nasal washes (A) and tissues (B) from guinea pigs intranasally inoculated with pH1N1 virus. The composition of the inoculum is shown on the far left of panel A. In (A), each barcode is represented by a colored box, the height of which indicates barcode frequency on a linear scale. The total height of the bar is scaled to the viral titer in log_10_ pfu/mL. In the pie charts shown in (B), each barcode is represented by a colored wedge, the size of which indicates barcode frequency. Pie charts are arrayed onto a schematic diagram of the respiratory tract according to the location of sampling. Shannon diversity is compared among the different regions in the respiratory tract (C). *p<0.05, ns: p>0.05 between indicated groups using Mann-Whitney-U or Kruskal-Wallis test with Bonferroni correction. The box represents the quartiles while the whiskers represent the upper and lower limits of the values.

Animals infected through aerosol and transmission-based exposures showed low barcode diversity overall in nasal wash and tissue samples (Figure 6-7). No significant differences in diversity were observed between URT and LRT, among lung lobe locations, or between proximal and distal sections within the lung lobes (p > 0.05 using Mann-Whitney-U or Kruskal-Wallis test with Bonferroni correction). Some patterns in barcode composition over space and time are of note, however, as outlined below.

**Figure 6.**
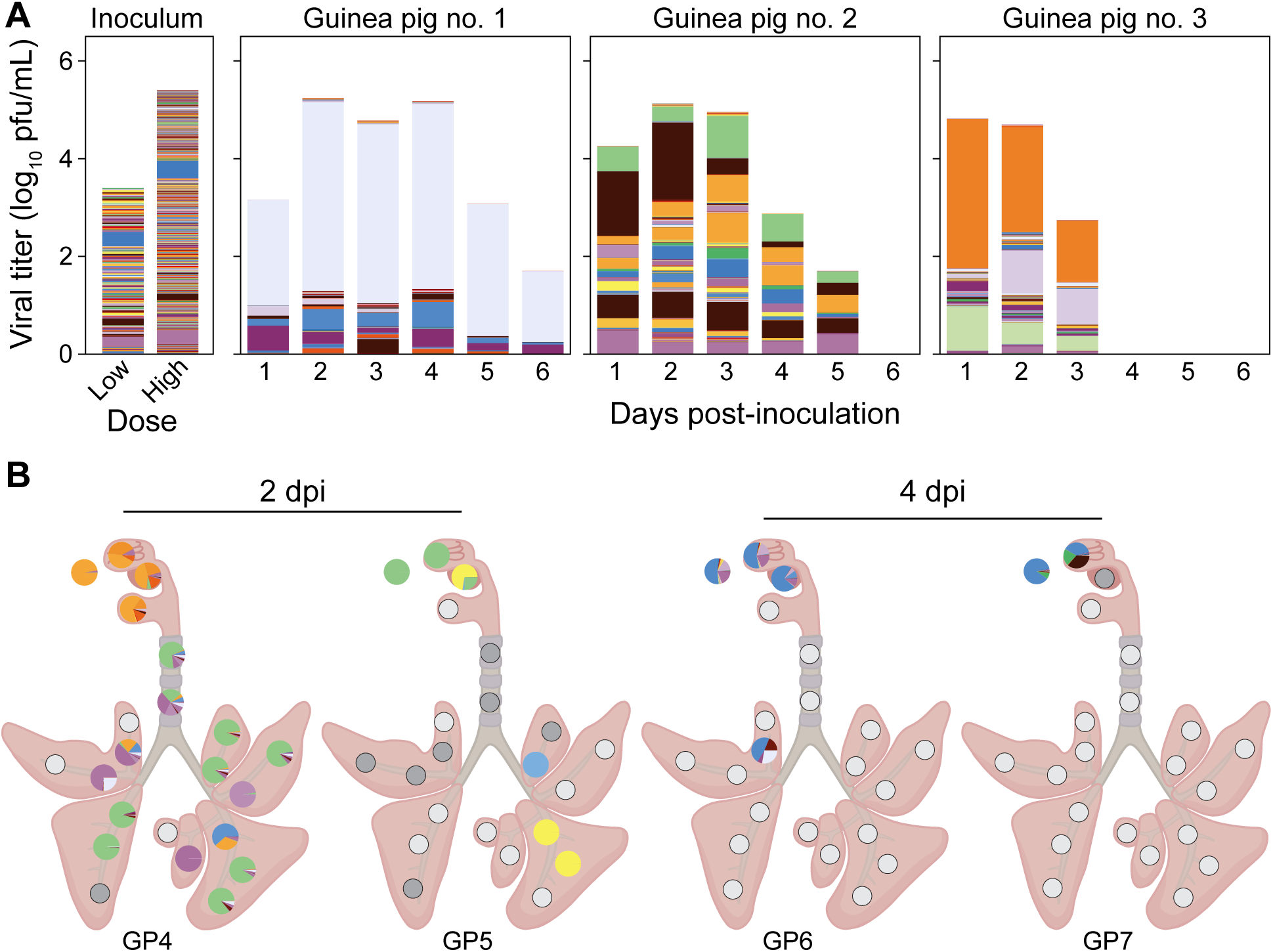
Barcode composition is associated with lung airways for animals exposed to pH1N1 aerosols. Viral barcodes detected in nasal washes (A) and tissues (B) from guinea pigs exposed to 1.53 x 10^2^ pfu of nebulized pH1N1 virus. In (A) each barcode is represented by a colored box, the height of which indicates barcode frequency on a linear scale. The total height of the bar is scaled to the viral titer in log_10_ pfu/mL. In the pie charts shown in (B), each barcode is represented by a colored wedge, the size of which indicates barcode frequency. Pie charts are arrayed onto a schematic diagram of the respiratory tract according to the location of sampling.

**Figure 7.**
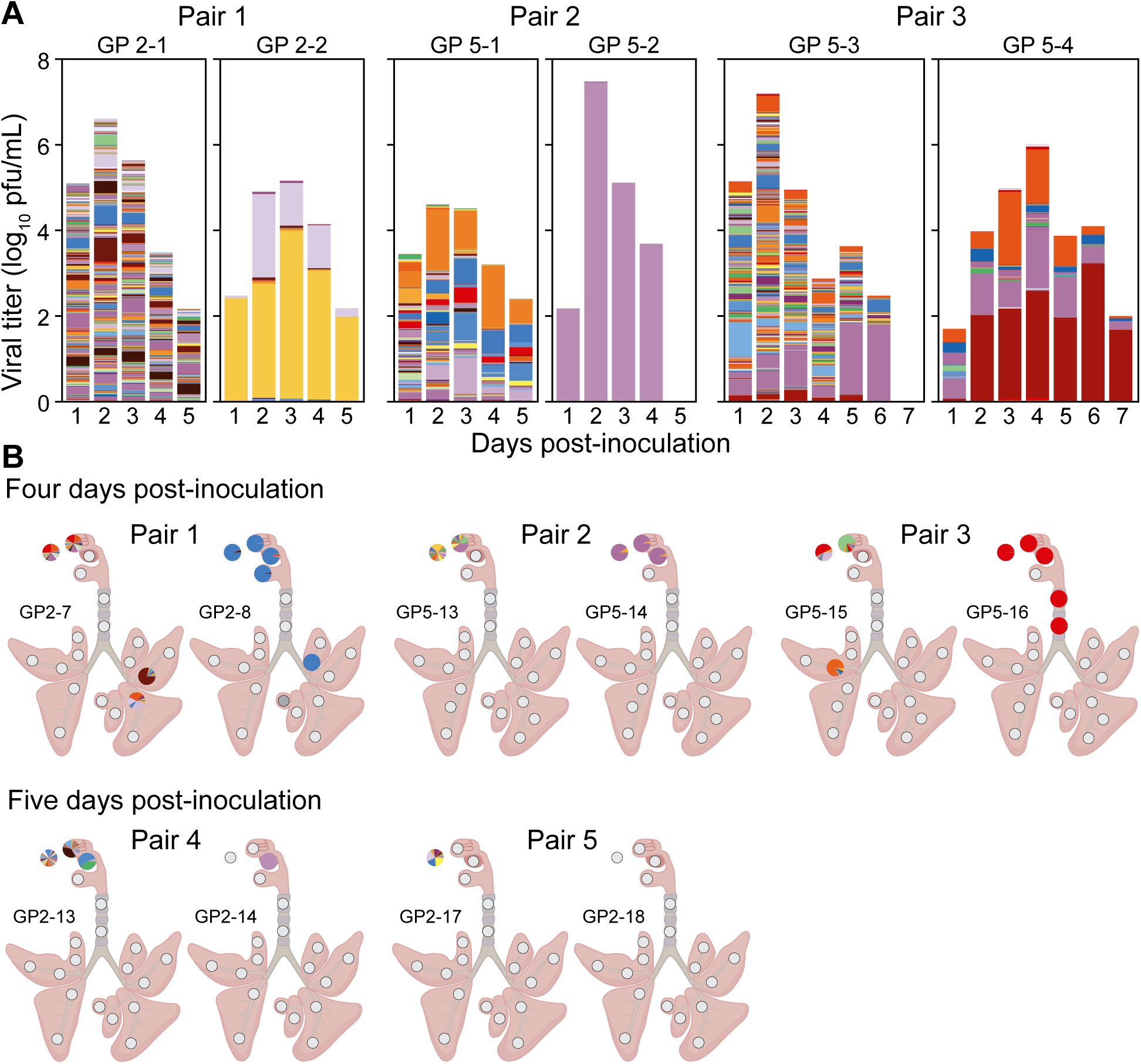
Bottlenecking at transmission is apparent in both upper and lower respiratory samples from recipient guinea pigs. Viral barcodes detected in nasal washes (A) and tissues (B) from donor and recipient guinea pigs. For each transmission pair, the donor animal is shown on the left and the recipient animal is shown on the right. In (A) each barcode is represented by a colored box, the height of which indicates barcode frequency on a linear scale. The total height of the bar is scaled to the viral titer in log_10_ pfu/mL. In the pie charts shown in (B), each barcode is represented by a colored wedge, the size of which indicates barcode frequency. Pie charts are arrayed onto a schematic diagram of the respiratory tract according to the location of sampling.

In aerosol inoculated animals, although diversity was low in the URT, it was stable over time (Figure 6A). Strong spatial heterogeneity between URT and LRT was, however, apparent at 2 dpi, potentially due to multiple independent seeding events at the time of inoculation (Figure 6B). At 4 dpi, LRT tissues were largely negative for virus, despite ongoing replication in the URT (Figure 6B).

In transmission experiments, donor guinea pigs inoculated intranasally showed relatively high barcode diversity in the URT, as expected. Barcode profiles in recipient animals, by contrast, were dominated by 1-3 barcodes regardless of the sample type and timepoint (Figure 7). Thus, the effects of a stringent transmission bottleneck were apparent. Of note, in the two recipient animals where viral positivity was detected in both URT and LRT (GP2-8 and GP5-16), similar barcode compositions were seen across these locations, suggesting they were populated by within-host dissemination following a single transmission event.

## Discussion

Influenza virus infection of the LRT is a major cause of morbidity and mortality (9–11). The features of such infections are not well-understood, however, due to the invasive nature of procedures necessary to obtain relevant specimens. To further understanding of influenza virus tropism and dynamics throughout the respiratory tract, we employed a guinea pig model. We first evaluated the spatial distribution of pH1N1 and seasonal H3N2 viral populations replicating within this host, finding that the former shows broader tropism, extending to the LRT. We then leveraged a barcode system to examine viral population dynamics, revealing that distance from the upper respiratory tract and airway branching impose genetic bottlenecks, such that pH1N1 populations replicating in the URT and different sites of LRT can be highly dissimilar.

Virus dose, mode of exposure, and inoculum volume have all been shown to modulate the course of influenza virus infection (31–34). Owing to direct delivery to the lungs, intratracheal and intranasal instillations with larger volumes often produce more severe disease (31, 34). The volume needed to reach the lungs is dependent on the host species: a 1 mL intranasal inoculum in ferrets and a 35 or 50 μL volume in mice are sufficient to induce similar outcomes (31, 34). In guinea pigs, a 300 μL intranasal inoculum is typically used to model influenza virus infection (35–38) and we show here that this volume reaches the lungs, whereas 30 or 100 μL volumes remain the nasal passages. We also note that some of the inoculum is deposited in the oral cavity and digestive tracts, although its impact on the course of influenza virus infection is unclear. Notably, viral detection in the lungs following intranasal inoculation was more robust than that seen following transmission or aerosol inoculation, despite comparable titers in nasal washes. This may suggest that seeding of the LRT is limited during transmission, although the timing of our sampling may have missed the window of opportunity to observe more widespread replication in the lungs. A longitudinal study in ferrets using a bioluminescent pH1N1 virus has shown that virus can be consistently detected in both the URT and LRT with direct virus inoculation, whereas heterogeneity in virus distribution in the URT and LRT was observed in recipient ferrets co-housed with a donor ferret (39). In this case, recipient animals either had virus in the URT only, LRT only, or both URT and LRT.

Prior work in guinea pigs has consistently shown robust replication in the URT with a range of seasonal influenza A and B viruses (25, 35, 36, 38, 40–42). Studies examining LRT involvement in this model are rare, but influenza A/HK/1/68 (H3N2) virus was found to yield histopositivity in the lungs when delivered intranasally at high dose (43). In ferrets, differential tropism between seasonal H3N2 and pH1N1 viruses consistent with that seen here has been documented (44–46). Also in humans, reports from the 2009 pandemic indicate heightened involvement of the LRT with pH1N1 infection compared to H3N2 or seasonal H1N1 infection (47).

IAV tropism is modulated by preferential binding to sialic acid receptors that vary in prevalence across different regions of the respiratory tract (48–50) and by the temperature sensitivity of viral replication (51, 52). Like seasonal IAV, isolates of the 2009 pandemic - such as the A/CA/07/2009 strain used here - bind alpha 2,6-linked sialic acid receptors (53, 54). Specificities within this category of sialyated glycans differ, however (55–57), and these differences may contribute to the broader tropism of pH1N1 viruses. In addition, while many seasonal IAVs show more robust replication at 33°C, a temperature mimicking the human URT, early pH1N1 viruses show comparable replicative potential at 33°C and 37°C (58, 59). Robust replication at 37°C is likely to be important for viral dissemination to the LRT.

By monitoring viral barcodes over time and at multiple locations throughout the respiratory tract, we can gain detailed insight into the genetic structure of viral populations as they disperse, expand, contract and transmit to contacts. Consistent with prior studies in ferrets that used similar approaches (16, 17), we saw that barcode diversity following intranasal inoculation is maintained in URT and trachea but lost upon dispersal to the lungs. We could furthermore detect patterns of reduced diversity with greater bronchial branching and, within a lobe, increased distance from major airways. These patterns suggest that within-host bottlenecking arises due to the physical loss of viral particles during transit through the airways, either at the time of inoculation or during secondary dispersal. Additional constrictions in diversity may follow from host antiviral responses active at the site of deposition and/or the heterogeneity of viral burst size at the single cell level (60–62), which would act during the establishment of infection at a new site.

The low barcode diversity that we observed in the URT following aerosol inoculation with pH1N1 contrasts with that seen in ferrets using a similar approach (16). While this difference may relate to host species or the differing instrumentation used to aerosolize the virus, the most likely contributing factor is the viral dose delivered. In ferrets, the aerosol exposure was designed to deliver 1×10^6^ PFU, while the estimated dose delivered to guinea pigs herein was 1.53×10^2^ PFU.

Low barcode diversity was also observed following transmission, consistent with the tight bottleneck described previously in ferrets, guinea pigs and naturally infected humans (30, 63–66). We reported recently that this tight bottleneck follows from losses of diversity at two stages: inter-host transfer and intra-host establishment (30). This two-stage process is not apparent in the data reported here, with even the earliest samples from recipients showing extremely low diversity. We attribute this difference to the transience of the more diverse viral population delivered to recipient animals, and the differing kinetics of viral establishment and expansion for different IAV strains (19, 40).

In summary, our data extend prior observations of the differing tropisms of H3N2 and pH1N1 viruses within guinea pigs. With examination of the population genetic changes accompanying pH1N1 spread throughout the respiratory tract, our data furthermore implicate long distance dispersal through the airways as a mechanism imposing stochastic losses of diversity. The spatial heterogeneity of the within-host viral population that results creates the potential for multiple divergent evolutionary paths to be followed within one host.

## Materials and Methods

### Barcoded viruses

Two viruses were used namely, influenza A/California/07/2009 (H1N1) virus (referred to as pH1N1 throughout) and influenza A/Texas/50/2012 (H3N2) virus (referred to as H3N2 throughout). Both viruses were generated through an eight-plasmid virus rescue system (67). For pH1N1, the neuraminidase gene was modified to become biallelic at 12 sites (5’ <u>R</u>CA TTC <u>M</u>AA TGG <u>R</u>AC CAT <u>W</u>AA <u>R</u>GA CAG <u>R</u>AG <u>Y</u>CC <u>W</u>TA TCG <u>R</u>AC <u>Y</u>CT AAT GAG CTG TCC <u>Y</u>AT <u>W</u> 3’) as described previously (16, 68). Modified DNA libraries containing biallelic sites were synthesized (Genewiz, South Plainfield, New Jersey) and cloned into parental plasmids using NEB Builder Hifi DNA Assembly (New England Biolabs, Ipswich, Massachusetts). Prior to assembly, parental plasmids were modified to contain an XhoI site and two stop codons to facilitate elimination of wild type sequence from the final virus preparations by (1) restriction digestion prior to DNA assembly and (2) by negative selection of truncated viral proteins during virus replication, respectively. Plaque assays on MDCK cells were used to quantify virus titer. Viruses were diluted in PBS according to the indicated dose to prepare inocula for all modes of exposure.

### Guinea pig experiments

Hartley guinea pigs purchased from Charles River Laboratories and weighing 300-350 g were used for all experiments. Each experimental group consisted of 1:1 ratio of female-male guinea pigs. For intranasal inoculations, nasal washes, and euthanasia, sedation was achieved by intramuscular injection of a ketamine-xylazine mix (9 mg and 1.24 mg per guinea pig respectively).

To determine the destination of liquid delivered intranasally, 30, 100, or 300 μL of blue tissue marking dye (Polysciences, Warrington, PA) was instilled intranasally. Guinea pigs were allowed to recover from sedation and proceed to normal activities for 30 minutes prior to euthanasia and tissue harvest.

For intranasal inoculations, each guinea pig received 300 μL containing barcoded pH1N1 or H3N2 virus at 5×10^4^ plaque forming units (pfu) by intranasal instillation. For transmission experiments, each donor guinea pig was similarly intranasally inoculated with 300 μL containing 5×10^4^ (first experiment) or 1×10^4^ pfu (second experiment). Recipient guinea pigs were then co-housed with donor guinea pigs after 24 hours.

For aerosol exposure, guinea pigs were placed into CODA Monitor Rat Holders (Kent Scientific, Connecticut, USA) that were modified by replacing removable parts with custom 3D-printed nose cone and rear plug. The nose cone was then connected to an Aerogen solo nebulizer (Aerogen, Illinois, USA) through a three-way T-piece splitter such that each nebulizer delivered virus to three guinea pigs at the same time. The guinea pigs were exposed to aerosols generated from a 600 μL volume containing either 5×10^6^ (high) or 5×10^4^ (low) pfu per guinea pig. The amount of virus delivered to each guinea pig is estimated to be 1.53×10^2^ (high) or 1.53 pfu (low) according to quantification procedure described below. The inoculum was placed into the nebulizer cup 75 μL at a time, allowing each droplet to completely nebulize before placing another one.

To avoid artificial dispersal of viral populations during nasal washing, nasal washes were performed in one set of guinea pigs while tissue harvests were performed in a distinct set of guinea pigs. Nasal washes were done by instillation of sterile phosphate buffered saline (PBS) drop-by-drop into each nostril and collection of refluxed fluid onto a sterile Petri dish.

All tissue sections were placed into tissue disruption tubes pre-filled with 1.7 mm Zirconium beads (OPS Diagnostics, Lebanon, NJ) and 1 mL PBS. Homogenization was done by bead beating samples twice for 1 minute at 6 m/s using a FastPrep-24 machine (MP Biomedicals, Solon, OH). Homogenates were clarified by centrifugation at 15,000 rcf for 5 minutes.

For lung sections, punches were taken using 6 mm biopsy punches (Integra™ Miltex®, Princeton, NJ). Tracheas were divided in half into upper and lower sections then cut into 2-4 rings to facilitate homogenization. Nasal turbinate tissues were harvested using a 5.5-inch Ruskin bone cutter (Roboz, Gaithersburg, MD) to the expose nasal cavity and a No. 12 scalpel to excise nasal epithelia.

Anterior nasal and oral swabs were collected post-mortem using micro ultrafine tip PurFlock Ultra® swabs (Puritan Medical Products, Guilford, ME) which was placed in a vial containing 1 mL brain-heart infusion broth (Sigma Aldrich, St. Louis, MO) supplemented with 100 IU/mL penicillin (Corning, Glendale, AZ), 100 μg/mL streptomycin (Corning), 200 μg/mL gentamicin (Gibco, Grand Island, NY), and 5 μg/mL amphotericin B (Gibco).

For histopathology and immunohistochemistry, tissues were fixed in 10% neutral-buffered formalin for a minimum of 24 hours. Lungs were inflated with 10% neutral-buffered formalin to preserve tissue architecture.

All experimental procedures were reviewed and approved by the Institutional Animal Care and Use Committee of Emory University.

### Quantification of virus delivered by aerosols

A group of 3 guinea pigs was exposed to aerosols as described above. To overcome the limit of detection of viral genomes, we used a higher initial inoculum of 5×10^6^ pfu in 600 μL per guinea pig. The guinea pigs were euthanized 1 hour post-inoculation and necropsy was performed. Nasal turbinates, soft palate, trachea (upper and lower sections) were harvested in 1 mL PBS and homogenized with tissue disruption tubes as described above. Entire lungs were cut in 1 cm cubes, placed in 3 mL PBS, and homogenized by mortar and pestle with the aid of sterile molecular grade sand (Fisher Scientific). Tissue homogenates were centrifuged for 5 minutes at 15,000 rcf and 140 μL of the supernatant was used for RNA extraction using the Qiamp Viral RNA Mini kit (Qiagen). To determine the pfu: RNA copy ratio of the pH1N1 virus, RNA from the virus stock was also extracted.

Copy number of viral RNA present in tissues and virus stock were quantified using droplet digital RT-PCR. Briefly, reverse transcription was done using the Maxima RT reagents (Thermo Fisher) with 1:1 mix of 6 μM UnivF(A)+6 and UnivF(G)+6 (5’-GCG CGC AGC AAA AGC AGG-3’ and 5’-GCG CGC AGC GAA AGC AGG-3’, respectively) at 55°C for 30 minutes, followed by heat inactivation at 85°C for 10 minutes. The cDNA products were diluted in 1:200 water and used as template for PCR with the QX200 ddPCR EvaGreen® Supermix (BioRad) and the primers NL_NP_309_F (5’-CCC TAA GAA AAC AGG AGG ACC C-3’) and NL_NP_411_R (5’-TTG GCG CCA AAC TCT CCT TA –3’). Droplets were generated using the QX200™ Droplet Generator (BioRad) and subsequently placed under the following thermocycling conditions: 95°C for 5 minutes; 40 cycles of 95°C for 30 seconds and 57.5°C for 1 minute; 4°C for 5 minutes; 90°C for 5 minutes; and a hold at 4°C. PCR positive droplets were quantified using the QX200™ Droplet Reader.

The RNA copy number quantified by droplet digital PCR was converted to infectious virus titer using the pfu: RNA copy number of the pH1N1 virus stock, determined empirically to be 17.4 copies/pfu.

### Immunohistochemistry

All staining procedures were performed at HistoWiz, Inc, using the Leica Bond RX automated stainer (Leica Microsystems) using a Standard Operating Procedure and fully automated workflow. Samples were processed, embedded in paraffin, and sectioned at 4 μm. Slides were dewaxed using xylene and alcohol based dewaxing solutions. Epitope retrieval was performed by heat-induced epitope retrieval (HIER) of the formalin-fixed, paraffin-embedded tissue using citrate based pH 6 solution (Leica Microsystems, AR9961) for 20 min at 100°C. Tissues were first incubated with peroxide block buffer (Leica Microsystems), followed by incubation with the rabbit Influenza A NP Polyclonal antibody (Invitrogen, PA5-32242) at 1:500 dilution for 30 min, followed by DAB rabbit secondary reagents: Polymer, DAB Refine and Hematoxylin (Bond Polymer Refine Detection Kit, Leica Microsystems) according to the manufacturer’s protocol. The slides were dried, coverslipped (TissueTek-Prisma Coverslipper) and visualized using a Leica Aperio AT2 slide scanner (Leica Microsystems) at 40X. Duplicate slides were stained with hematoxylin and eosin for histopathology.

Immunostaining summary score refers to the overall subjective semi-quantitative evaluation of intensity and number of positive cells across all compartments of the tissue examined (69–71). The scores are given as follows: 0 for no staining, 1 for small amount with <10% positivity, 2 for mild-moderate amount with 11-20% positivity, 3 for moderate amount with 21-40% positivity, 4 for moderate-large amount with 41-60% positivity, and 5 for large amount with >60% positivity.

### Amplicon sequencing

RNA extraction from nasal washes and tissue homogenates was done using the NucleoMag RNA extraction kit (Machery-Nagel, Düren, Germany) and EpMotion liquid handler (Eppendorf, Hamburg, Germany) according to the manufacturer’s protocol. The barcoded region of pH1N1 was amplified using the Superscript III One-step RT-PCR with Platinum Taq High Fidelity DNA polymerase (Thermo Fisher) and the following primers: 5’-TCG TCG GCA GCG TC AGA TGT GTA TAA GAG ACA G<u>AC CTT CTT CTT GAC TCA AGG G</u>-3’ (forward) and 5’-GTC TCG TGG GCT CGG AGA TGT GTA TA AGA GAC AG<u>C AAG CGA CTG ACT CAA ATC TTG A</u>-3’ (reverse). Influenza-specific regions are underlined. The rest of the primer sequence comprises of Illumina adapters that allow for direct indexing. Thermocycling conditions were: 55°C for 2 minutes; 45°C for 60 minutes; 94°C for 2 minutes; 30 cycles of 94°C for 30 seconds, 59°C for 30 seconds, 68°C for 1 minute; final extension at 68°C for 5 minutes, and hold at 10°C. For all DNA preparations, DNA fragments were purified using NucleoMag NGS Clean-up and Size Select (Macherey-Nagel) at a 2x (v/v) bead to DNA preparation ratio. Quantification of purified DNA was done using the Qubit dsDNA high sensitivity kit (Thermo Fisher). Libraries were prepared by amplifying RT-PCR products with Nextera XT Index kit v2 set A indexes (Illumina, San Diego, California) and KAPA HiFi HotStart ReadyMix (Roche, Basel, Switzerland). After another round of DNA purification, the libraries were manually pooled at equimolar concentrations and run using a Miseq v2 kit with 2×150 bp or 2×250bp output (Illumina). The Emory Integrated Genomics Core performed all sequencing on Illumina Miseq instruments.

### Amplicon analysis

Barcode frequencies and diversity metrics were determined using BarcodeID (https://github.com/Lowen-Lab/BarcodeID) as previously described (16, 30).

## Data Availability

Sequences are available through NCBI’s Short Read Archive (https://www.ncbi.nlm.nih.gov/sra) BioProject accession number xxxxxxxxx (pending at the time of submission). All custom computer code necessary to reproduce the results presented in the manuscript are available on GitHub. The pipeline for the analysis of barcode composition is at (https://github.com/Lowen-Lab/BarcodeID).

## Acknowledgments

This study was supported in part by the Emory Integrated Genomics Core (EIGC; RRID:SCR_023529), which is subsidized by the Emory University School of Medicine and is one of the Emory Integrated Core Facilities. Additional support was provided by the Georgia Clinical & Translational Science Alliance of the National Institutes of Health under Award Number UL1TR002378. The content is solely the responsibility of the authors and does not necessarily reflect the official views of the National Institutes of Health. The research was funded by NIH/NIAID through R01 AI 165644, R01 AI 154894 and the NIAID Centers of Excellence for Influenza Research and Response (CEIRR), contract number 75N93021C00017 to ACL.

